# Significance estimation for large scale untargeted metabolomics annotations

**DOI:** 10.1101/109389

**Authors:** Kerstin Scheubert, Franziska Hufsky, Daniel Petras, Mingxun Wang, Louis-Félix Nothias, Kai Dührkop, Nuno Bandeira, Pieter C Dorrestein, Sebastian Bocker

## Abstract

The annotation of small molecules in untargeted mass spectrometry relies on the matching of fragment spectra to reference library spectra. While various spectrum-spectrum match scores exist, the field lacks statistical methods for estimating the false discovery rates (FDR) of these annotations. We present empirical Bayes and target-decoy based methods to estimate the false discovery rate. Relying on estimations of false discovery rates, we explore the effect of different spectrum-spectrum match criteria on the number and the nature of the molecules annotated. We show that the spectral matching settings needs to be adjusted for each project. By adjusting the scoring parameters and thresholds, the number of annotations rose, on average, by +139% (ranging from −92% up to +5705%) when compared to a default parameter set available at GNPS. The FDR estimation methods presented will enable a user to define the scoring criteria for large scale analysis of untargeted small molecule data that has been essential in the advancement of large scale proteomics, transcriptomics, and genomics science.

## Introduction

Untargeted mass spectrometric analysis of small molecules is important in our understanding of all molecules in the environment, ocean, and individual organisms.^1–4^ In untargeted mass spectrometry experiments, tandem MS (MS/MS) spectra are collected of molecules present in the analytical sample. To annotate these unknowns, the MS/MS spectra are compared against a library of reference MS/MS spectra.^5–8^ At present, spectrum-spectrum matches of unknown and library spectra are scored but this score alone provides no statement about statistical accuracy of that assignment. Without statistical techniques in place to estimate false discovery rates of identifications, researchers do not have a guide to set appropriate scoring criteria, unlike proteomics, peptidic small molecule identification, transcriptomics and genomics where statistical assessment and false discovery calculations for annotations are the norm.^9–12^ This leads untargeted liquid chromatography tandem mass spectrometry (LC-MS/MS) based metabolomics or any other small molecule based untargeted mass spectrometry analysis to yield identification results where errors rates are uncontrolled that can lead to lack of sensitivity or worse, rampant false discoveries.

To compound the challenge, due to advances in instrumentation and the re-emergence of appreciation in the function of small molecules, the scientific community is generating more and more untargeted mass spectrometry data. These LC-MS/MS based experiments are now commonly applied in medicine, life science, agriculture toxicology, exposome, ocean and forensic research to name a few. Hundreds to thousands of MS/MS spectra are generated from a single sample with modern instruments and collectively tens of millions of MS/MS spectra for large scale projects. There are also a growing number of MS/MS spectra available in public spectral libraries.^5–7,13,14^ To most in the scientific community, including mass spectrometrists and metabolomics investigators themselves, it often comes as a surprise that there is no significance estimation in metabolomics annotations yet, like it has been adopted, and thereby advanced, the fields of proteomics, genomics and transcriptomics. While guidelines and rules have been established for reporting the annotation of molecules results from MS-based metabolomics data by the metabolomics standards initiative^15^, or golden rules^16^, these are not commonly reported in the majority of metabolomics studies, are subject to interpretation, and need to be updated regularly to reflect current scientific capabilities and advances. Manual validation at the scale of tens of thousands to millions of spectra library matches is not realistic to do for each large scale experiment, and automated solutions for the annotations that enable downstream analysis such as pathway mapping, metabolism analysis and ultimately prioritization for manual validation are needed; but this process starts with annotations. Different from *in silico* annotation of MS/MS spectra of peptidic small molecules^17^, our methods for building decoy spectral libraries do not generate artificial metabolite structures: both, the construction of metabolite structures which are plausible but non-existing in nature, and the prediction of fragmentation spectra from metabolites structures are extremely challenging problems^18,19^.

Therefore, we present and evaluate four different approaches to estimate the accuracy of the results. These four methods are empirical Bayes, which uses a probabilistic model of score distributions, and three target-decoy approaches where the decoy libraries are generated via the naive method, a spectrum-based and a fragmentation tree-based method. Using three test reference databases, Agilent^20^, MassBank^7^, and GNPS^6^, with thousands of MS/MS spectra that have the structures of molecules associated with them, we show that all but the naive methods can estimate false discovery rates^21^ (FDR, the proportion of false discoveries among the discoveries) and q-values (the minimal FDR thresholds at which given discoveries should be accepted) with high accuracy; also, that the best performing method, the fragmentation tree-based approach, can be used for FDR estimation at the scale of 10,000s of LC-MS/MS runs. The FDR estimation has now been implemented as a tool called `passatutto`, named after a food mill used to remove unwanted particles commonly used in Italian kitchens, and has been integrated into GNPS web-platform (http://gnps.ucsd.edu)^6^. `passatutto` provides experimentalists with a measure of confidence in MS/MS-based annotations by reporting an FDR, to guide the selection of scoring parameters for a project compatible with large scale MS-metabolomics projects. To validate the FDR approach and how it performs for spectral annotation with real large scale untargeted mass spectrometry, we performed FDR controlled spectrum library matching with 70 datasets from GNPS, consisting of thousands of LC-MS runs. Overall change in annotation rate was at +139%, ranging from −92% up to +5705% when compared to a conservative default scoring thresholds used for living data in GNPS. Further, with the given default scoring scheme we observed a range of FDRs between 0.0 and 23.7%, indicating that there exist no universal scoring criteria that can control the FDR in all datasets. This adaptive approach shows promise to both increase identifications and curb false positives in large scale metabolomics experiments.

## Results and discussion

Large scale non-targeted LC-MS/MS experiments result in hundreds to thousands of query spectra from a single chromatographic run. For molecular annotation these MS/MS spectra are typically searched against a spectral library, which in turn, results in spectral library hits that are sorted by score. Using a decoy spectral library to estimate FDR is common in proteomics; there, the decoy database is often a (pseudo-)reverse peptide database or a shuffled database^9,22,23^. This is possible for peptides and proteins because these are linear polymer chains over 23 proteinogenic amino acids, of which 20 amino acids are most common. The reason why target-decoy approaches for FDR estimation have not been applied so far to metabolomics, are the difficulties in generating decoy libraries; small molecules are diverse in structure, and shuffling or reversing a database is not possible. Therefore, alternative strategies needed to be developed for FDR estimation. Our first method uses an empirical Bayes approach^24^ whereas the second, third and fourth FDR estimation methods rely on the target-decoy approach, using different decoy databases (Figure 1a-d). Although the generation of “random” MS/MS spectra for small molecules is conceptually more challenging than for peptides^25^, it became possible with recent methodological advances^6,26–28^. To estimate the FDR using a decoy database, three strategies were devised to create the decoy MS/MS library (Figure 1b-d), where the first two methods are spectrum-based while the third is fragmentation tree-based^23,24^. To show compatibility with different spectral matching scoring schemes, we present results for the MassBank scoring and the GNPS scoring, both of which utilize modified versions of the cosine similarity (also known as normalized dot product). In principle, other commonly used scoring schemes for spectral matching, such as cosine similarity itself^28,29^, scorings based on the number of matching fragment ions and the sum of intensity differences^30^ or scorings which incorporate mass differences^31^, could be employed as well.

The key considerations that went into the design of the decoy spectral libraries was to ensure that decoy spectra mimic real spectra as closely as possible, but at the same time, do not correspond to MS/MS spectra of any true metabolites present in the sample. This ensures that hits in the decoy database are equally likely as false hits in the spectral library (the target database). In addition, we assured that for any precursor mass range, the same number of target and decoy spectra were found. All methods circumvent generating decoy structures, as it is unsolved problem to generate molecular structures which are sufficiently similar to the structures in the target spectral library, but not present in the sample. Generating decoy MS/MS spectra completely at random, i.e., randomly drawing both masses and intensities of the fragment ions, will not result in an adequate decoy spectral library, as there are ion masses that can be generated but will never be found in a real MS/MS spectrum. Addition of adducts to the spectra that are not encountered would be a solution to creating a decoy spectral library, as was recently done for parent mass FDR calculations for imaging mass spectrometry data^32^; but these adducts would not look like spectra that we would encounter in an MS/MS spectrum from a biological sample and therefore this solution is not appropriate for the annotation of MS/MS spectra.

For the *naive* decoy spectral library, we use all possible fragment ions from the reference library of spectra and then randomly add these ions to the decoy spectral library, until each decoy spectrum reaches the desired number of fragment ions that mimics the corresponding library spectrum (Figure 1b). This method is presented as a baseline evaluation of the other, more intricate methods. The second method is similar to the naive method, as we create the decoy spectral library through choosing fragment ions that co-appear in the spectra from the target spectral library (Figure 1c): In this *spectrum-based approach*, we start with an empty set of fragment ion candidates. First, the precursor fragment ion of the target spectrum is added to the decoy spectrum. For each fragment ion added to the decoy spectrum, we choose all spectra from the target spectral library which contain this fragment ion, within a mass range of 5 ppm. From these spectra, we uniformly draw (all fragment ions have the same probability to be drawn) five fragment ions that are added to the fragment ion candidate set; we use all fragment ions in case there are fewer than five. We draw a fragment ion from the fragment ion candidate set and add it to the decoy spectrum, then proceed as described above until we reach the desired number of fragment ions that mimics the corresponding library spectrum. Fragment ions with mass close (5 ppm) to a previously added fragment ion mass, or masses above the precursor fragment ion mass are discarded. If the precursor ion is absent from the MS/MS spectrum, we use the selected ion mass to find matching compound masses. The third solution is a *fragmentation tree-based* approach, where decoy spectra are generated using a re-rooted fragmentation tree (Figure 1d). From the original fragmentation tree, its structure and all losses are kept, and some new internal node is selected as new root, with the molecular formula of the precursor ion. Molecular formulas of all fragment ions are calculated along the edges of the tree, subtracting losses. In case the tree rearrangement yields chemically impossible molecular formulas (that is, a negative number of atoms for some element), the corresponding loss and its subtree are placed to another branch of the tree (re-grafted), attaching it to a uniformly selected node. The new root node is not drawn uniformly: Instead, a node is chosen as new root with relative probability 1/(n + 1), where n is the number of edges that we would have to re-graft. For all three methods, intensities of the original fragment ions are used.

**Figure 1.**
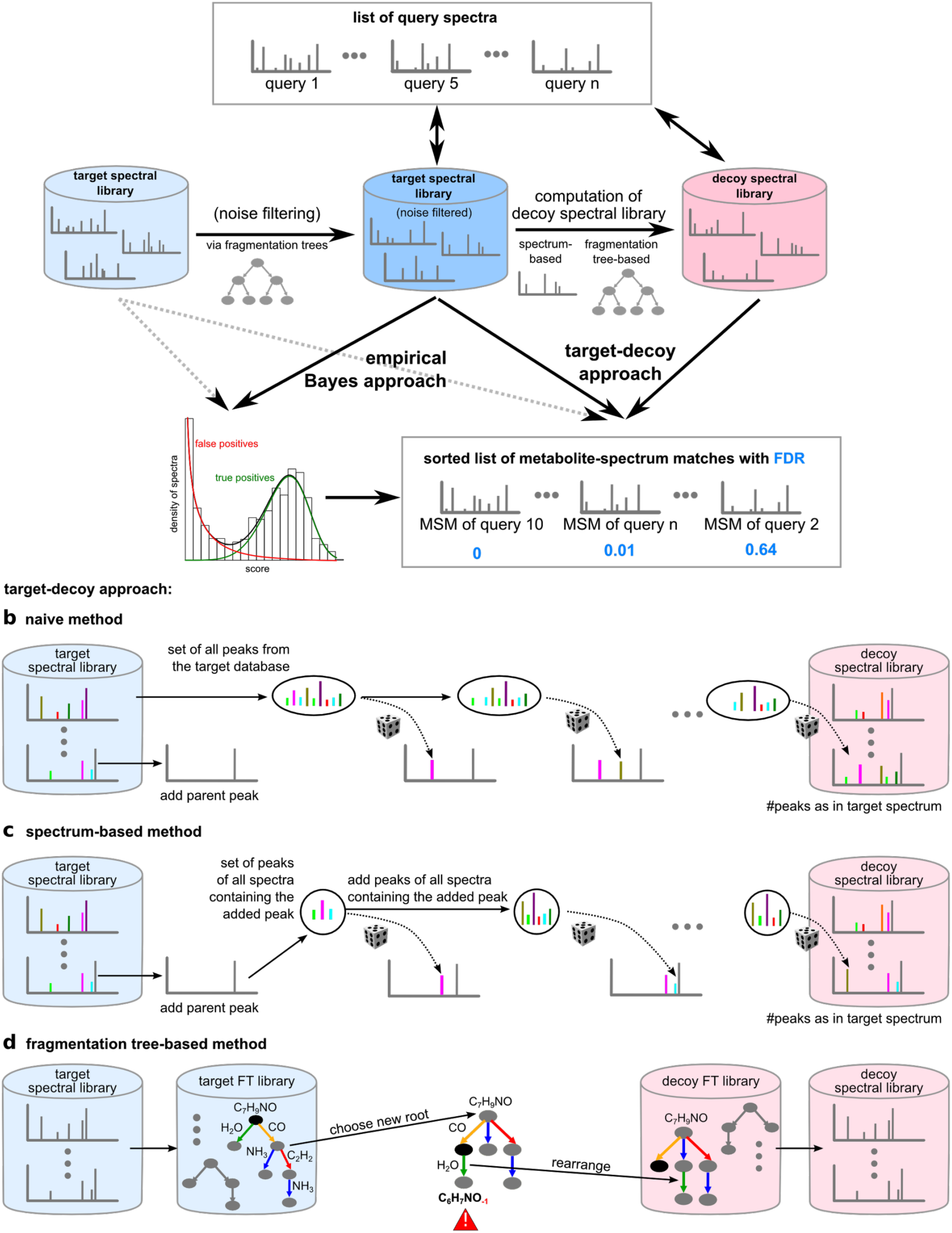
False discovery rate estimation. (a) Overview. The empirical Bayes approach estimates FDRs from a two-component mixture of distributions representing true and false hits (positive identifications). In the target-decoy approach, query spectra are searched in target and decoy spectral library, and FDRs are estimated from the merged and sorted list of spectrum matches. (b-d) To construct a decoy spectral library, we implemented three methods. (b) Naive method: Randomly adding fragment ions from the reference library to the decoy spectrum. (c) Spectrum-based method: fragment ions are iteratively added to the decoy spectrum, conditional on fragment ions that have previously been added. (d) Fragmentation tree-based method: A fragmentation tree is computed from the target spectrum, its root is relocated. New formulas of fragments are calculated according to the losses in the tree. Fragments with invalid formulas are relocated.

Assessing the quality of empirical Bayes, and the naive, spectrum-based and fragmentation tree-based target decoy databases was done by p-value estimation, and by testing q-value estimates against exact values using public MS/MS libraries. Evaluation can only be carried out when the true identity of all query compounds is known. To assess quality, we used high resolution reference spectra from the Agilent, MassBank and GNPS libraries. Only spectra that had the unfiltered spectrum in the public domain, that had SMILES or InChI structure annotations (line notations for describing chemical structure using short strings) and for which the parent mass matched to the exact structure-based mass to within 10 ppm, were used for the assessment of the FDR estimations. As an initial test, we checked if p-values of false hits (false positive identifications) estimated by our methods are uniformly distributed^33^: The p-value of a spectrum match is the probability to randomly draw a result of this or better quality, under the null hypothesis for which a spectrum has been randomly generated. We observe an almost uniform distribution of p-values, both for the empirical Bayes approach and the fragmentation tree-based target-decoy approach (Figure 2a-f). This agrees with the distribution of p-values under the null hypothesis, and shows that our decoy databases are indeed representative models of the null hypothesis.

**Figure 2.**
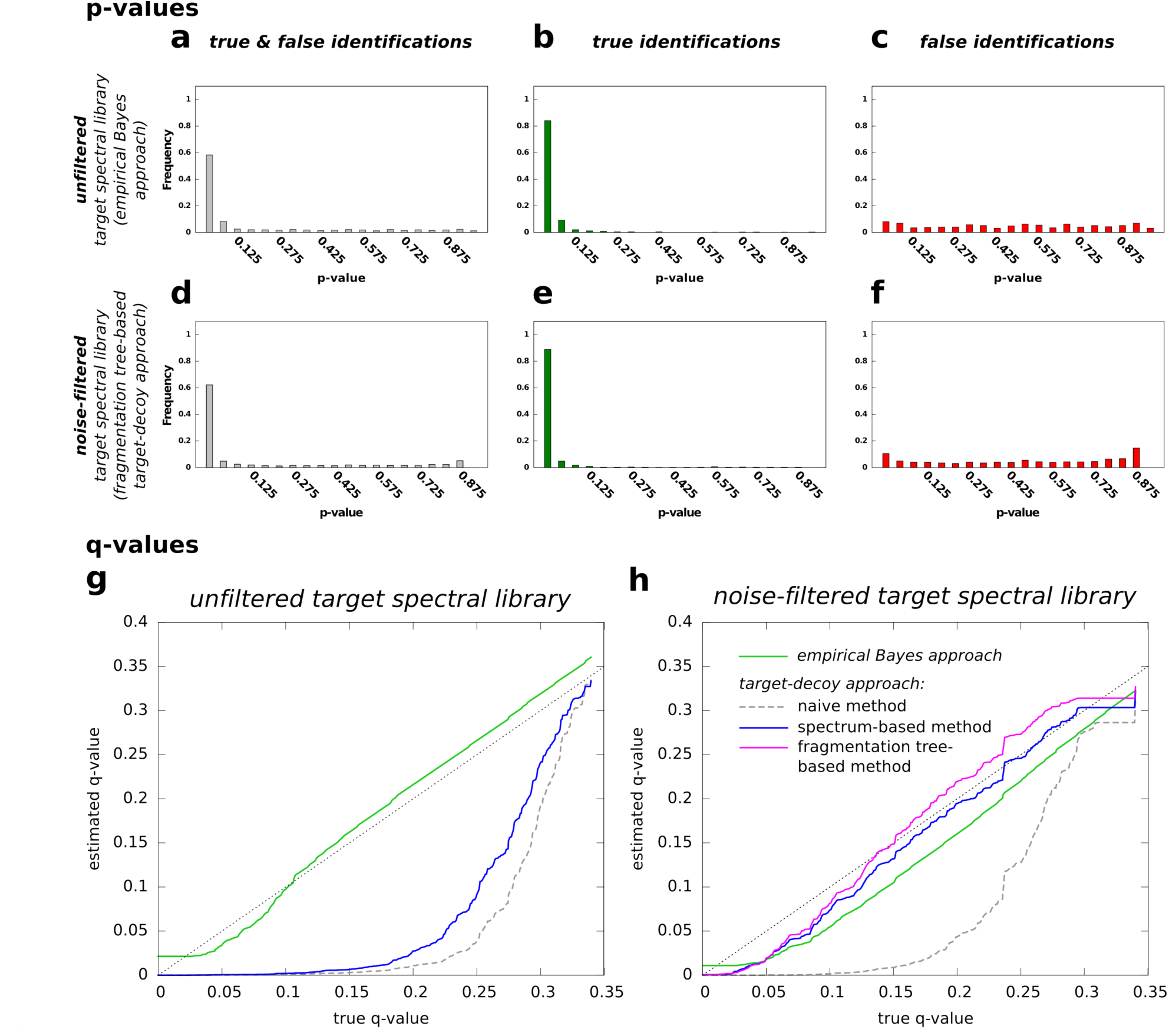
Quality assessment for FDR estimations. (a-f) p-values. Distribution of p-values for searching for Agilent query spectra to the GNPS library using the MassBank scoring function. For searching in the unfiltered target spectral library (a-c), p-values are calculated using the empirical Bayes approach. For searching the noise-filtered target spectral library, p-values are calculated using the fragmentation tree-based target-decoy approach (d-f). Distributions contain p-values from ten decoy spectral libraries. p-value distribution for both, true and false hits (a,d), p-value distribution for true hits only (b,e), and for false hits only (c,f). By definition, the distribution of p-values for false hits has to be uniform. (g,h) q-values. Estimated (y-axis) vs. true q-values (x-axis) for searching Agilent query spectra collected on a Q-tof in the unfiltered (g) and noise-filtered (h) version of the GNPS library using the MassBank scoring function. Results for empirical Bayes approach and the three target-decoy approaches. For the fragmentation tree-based method, we searched against the noise-filtered GNPS only, since this approach applies noise-filtering by design. The naive target-decoy approach can be seen as baseline method for comparison. For target-decoy methods, results are averaged over ten decoy spectral libraries.

To evaluate the quality of estimated FDRs, we compared q-values of the four methods presented here with true q-values; we are not aware of other methods for this estimation. In addition, we also assessed the impact of noise filtering on the quality of FDR estimation: Noise-filtering by fragmentation trees is accomplished by calculating a fragmentation tree that annotates some of the hypothetical fragment ions with molecular formulas^27,34^; only these annotated fragment ions are kept, resulting in a cleaned spectrum that only keeps fragment ions that are well-supported by the fragmentation process. For the unfiltered target spectral library, empirical Bayes approach resulted in good estimates, whereas spectrum-based target decoy did not work as accurately (Figure 2g, h): the empirical Bayes approach represented a good fit of the bisecting line, while the spectrum-based approach did not. For the noise-filtered target spectral library, the target-decoy methods except the naive method allow for accurate q-value estimates, but the fragmentation tree-based method performed slightly better (Figure 2c). The naive method never results in accurate q-value estimates: Even for true q-values around 0.15, estimates are already close to 0. All methods tend to overestimate significance; in particular, estimates are close to zero for true q-values below 0.05, in agreement with what has been previously observed in metaproteomics^35^.

To further evaluate the robustness of our estimates, we generate ten decoy spectral libraries for each decoy method. Because generating decoy spectral libraries is a random process, q-values vary slightly between the ten decoy spectral libraries; we found these variations to be negligible. Results in Fig. 2 present searching Q-TOF spectra using the MassBank scoring function. Results for the cosine similarity score were similar, showing applicability for different scoring approaches. Furthermore, using Orbitrap MassBank spectra as queries yielded similar results, indicating that the methods work even when the target spectral library is measured with different type of MS instruments.

We evaluated the fragmentation tree-based decoy FDR estimation method broadly across 70 datasets at GNPS. These datasets included high resolution Q-TOF or Orbitrap data from 6,220 LC/MS runs encompassing human, microbe, plant and marine-organism derived samples. To calculate both the 1% FDR and 5% FDR, the total running time for the FDR computation of the spectral library matches associated with all the projects took approximately forty-eight hours on the GNPS cluster, demonstrating the compatibility of the FDR approach with large-scale metabolomics experiments. At 1% FDR, the average gain in annotation for the 70 public data sets was 139% with a range of −92% up to 5705% in annotations when comparing the number spectral library hits retrieved from living data in GNPS that has a default cosine score of 0.7 (Figure 3). At a score of 0.7, the annotations from continuous identification, as judged by the community via a four-star rating of the identifications, the GNPS community provided feedback that 91% of the annotations are correct, 4% possible isomers or correct, 4% not enough information to tell and 1% is incorrect^6^. When using 5% FDR, a mean gain annotation of 235% was obtained and had a range of −75% up to 6705% gain (Figure 3). This result shows that the same spectrum matching score can contribute to a highly variable FDR and that the FDR can be drastically different for each project. This means that the spectral scoring for annotations needs to be adjusted on a per project basis and based on the false discovery rates the end user is willing to accept. With the 70 projects analyzed there were no trends with respect to the instrument type observed, in agreement with our benchmark results (Figure 2).

**Figure 3.**
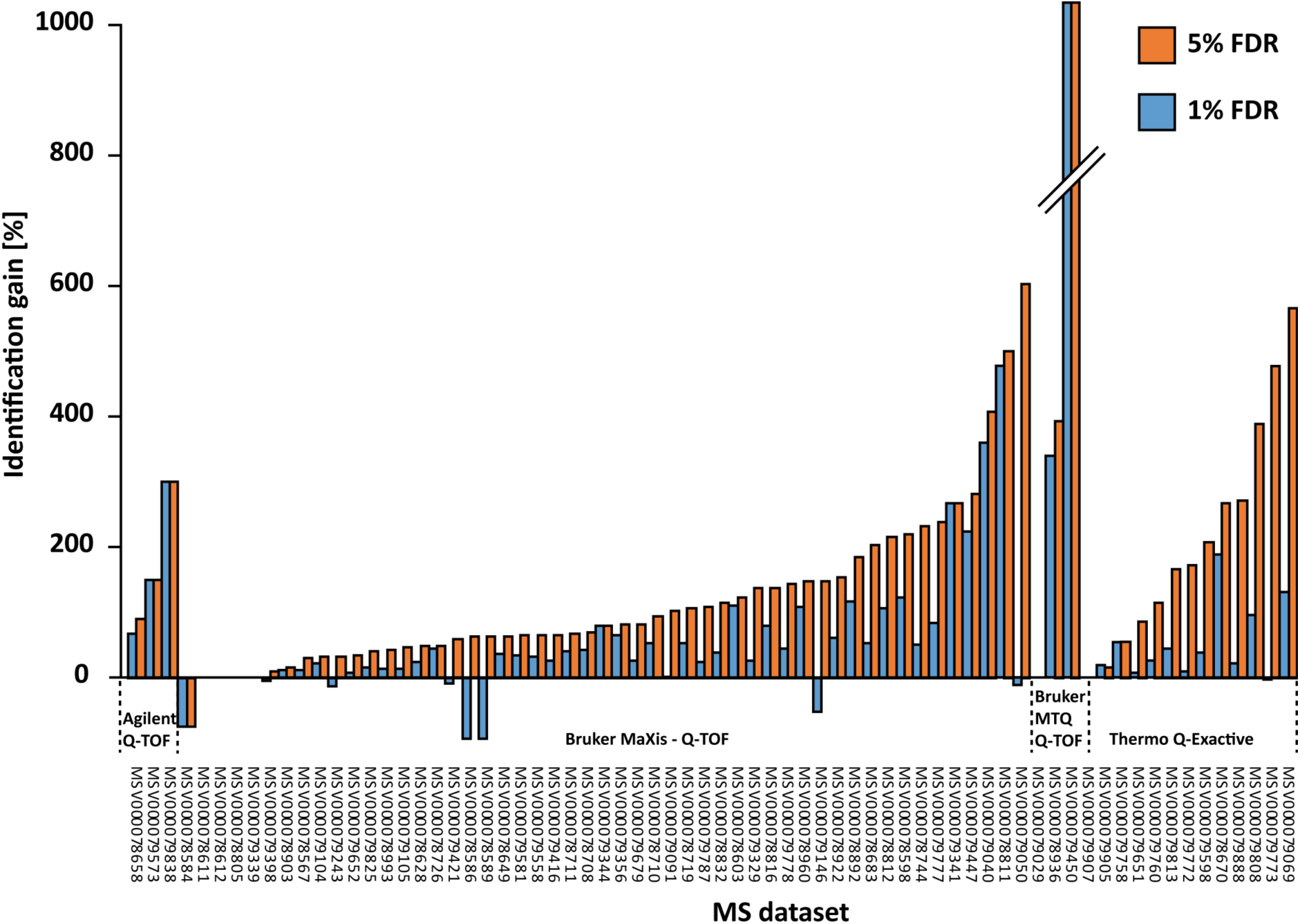
FDR based annotations for metabolomics 70 projects from human, microbes, plants, and marine-organism derived samples. The plot shows the percent gain in annotations for each of the data sets in GNPS-MassIVE at 1% and 5% FDR in relationship to the mass spectrometer used.

Further, we explore the impact of cosine scoring and the minimum number of fragment ions to match on the number of matches associated with 1% FDR using the fragmentation tree-based decoy strategy. Over the 70 public metabolomics projects, the minimum matched peaks was modulated resulting in a cosine threshold ranging from 0.3 to 1 with the number of identifications represented in a histogram (Figure 4a). The results reveal that the more ions that were required to match, the more forgiving the spectral scoring could be. When 8 ions were required to match, the most common score to achieve 1% FDR was found to be between a cosine of 0.50-0.60, while when two ions were required to match, the most common score required was 0.85-0.95, however nearly 40% of projects when 2 ions were required could not match a single spectrum in the database or the decoy and reveals an inherent limitation in untargeted mass spectrometry. For all the projects that require a cosine of 1 to achieve 1% FDR, not a single annotation was obtained (Figure 4b). We observed that the most number of annotations was achieved with a minimum of 5 fragment ions matching, with 6 ions and 4 ions as close second and third in terms of the number of spectra that were annotated. Interestingly as the number of fragment ions required to match the number of matches dropped to 3 and 2, the number of total matches decreased significantly. At these scores, there is not enough spectral information to differentiate a match to the library from the decoy library and therefore drives up the FDR quicker. However, there is a clear optimum because as we require 7 and 8 ions to match, we again see decrease in the number of annotations. This is because there are fewer spectra that have a minimum of seven or eight fragment ions to match.

**Figure 4.**
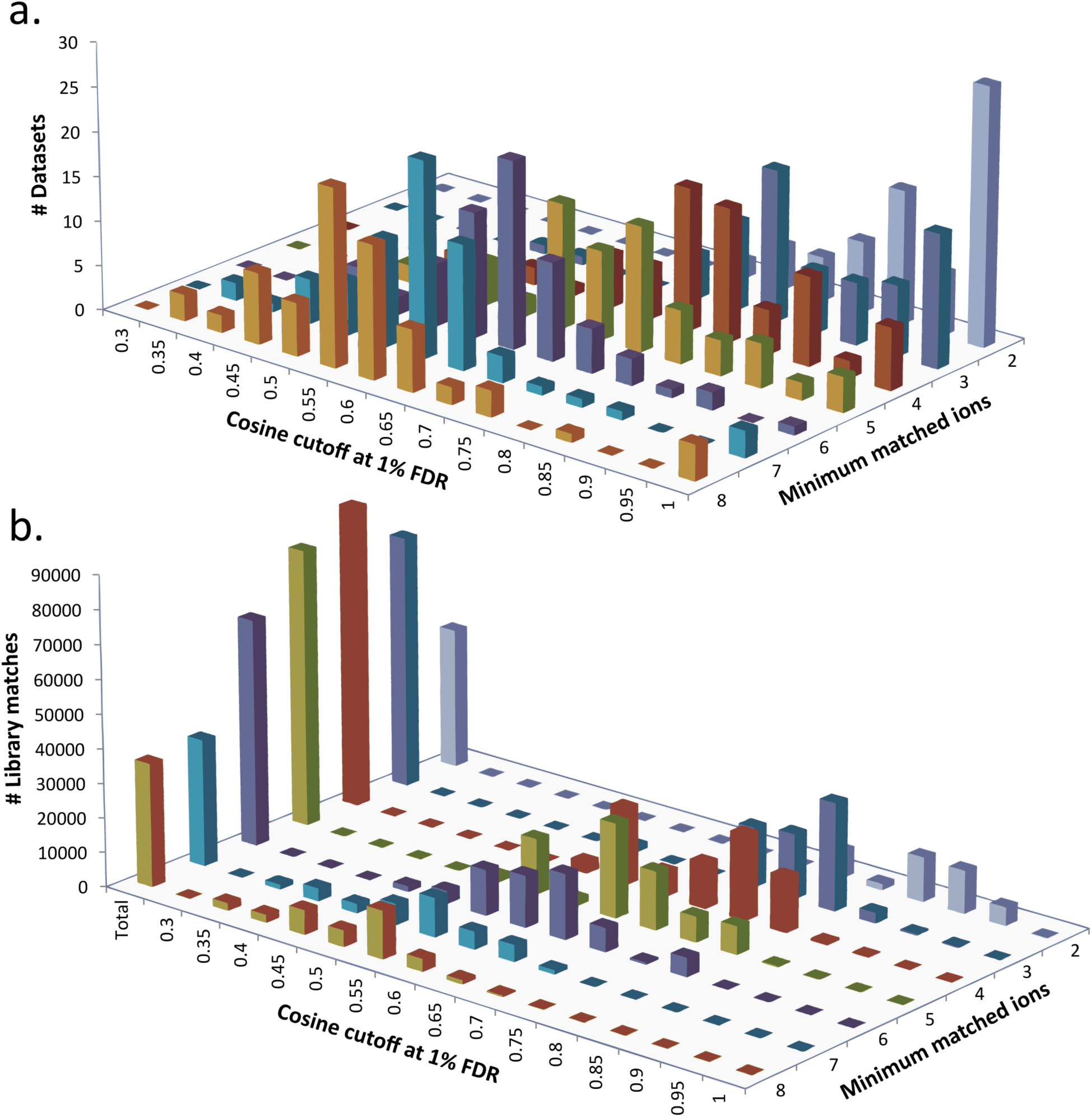
The impact of number of matching fragment ions in a spectrum and cosine score at 1% FDR. a. Frequency of datasets in relationship to number of minimum fragment ions to match and cosine at 1% FDR estimation. b. The number of MS/MS matches in relation to minimum matched fragment ions and cosine.

We can compare the results to the results from the default GNPS living scoring value of cosine of >0.7 and a minimum of 6 fragment ions to match^6^. This GNPS community assessed matches are the only direct comparisons we can currently make in the metabolomics field how the FDR estimation impacts results. This comparison revealed that for most of the projects, an FDR of 1% was achieved at cosine of 0.6-0.65 (for 5% FDR, most of the projects dropped to a cosine of 0.5-0.55) and therefore living data in GNPS is slightly more stringent. However, a key observation is that the data shows that for each data set the cosine scoring needs to be adjusted when compared to the default GNPS parameters of cosine of 0.7 or greater and minimum 6 fragment ions to match used with living data as we both an increase and decrease in the number of annotations. In other words, GNPS living data enabled through continuous identification^6^ that uses just one specific scoring value not only underestimates the annotations for most projects but perhaps more importantly, as this affects the interpretation of the results, living data also overestimates the number of annotations for some projects. At 1% FDR, 13% of the projects revealed that GNPS living data parameters overestimates the number of annotations and underestimated the annotations in 82% of the projects and the remaining 5% of projects remained unchanged by the introduction of the FDR estimation. There are, however, many molecules that do not provide 6 ions when fragmented. These are currently missed by living data in GNPS. Thus, FDR calculations enables an informed decision in terms of the analysis parameters that a researcher can use in terms of deciding what the level of acceptable incorrect annotations that can be expected with such parameters. These results demonstrated why the introduction of significance estimation and FDR assessments are critical for the field of untargeted small molecule mass spectrometry and that significance estimations needs to become a routine part of this field.

## Conclusion

There is more and more untargeted small molecule mass spectrometry research done by the scientific community. In addition, due to MS instrument advances the number of samples analyzed is also increasing and yet we are still using artisanal methods for assessing annotations and do not report the confidence in the annotations. This is remarkable because the interpretation of these data depends on annotations. We currently also do not have ways to assess the landscape of possible experimental analysis parameters that provide the annotations that is appropriate to use. Of the four FDR methods assessed, the fragmentation tree-based decoy strategy worked most effectively on noise filtered data. Our methods require that target mass spectra are noise-filtered. We use fragmentation trees^26,27^ to separate signal fragment ions from noise fragment ions to “clean" target spectra from spectral libraries. We demonstrated that this approach can be used for providing confidence measures in large scale metabolomics project, where it is becoming more and more impossible to inspect each annotation by hand, which is the current norm in metabolomics. It revealed that the spectrum scoring parameters need to be adjusted on a per-project-basis, which requires a form of confidence measures associated with the results. Although such evaluations have been critically important for advancing other fields such as proteomics, genomics and other fields, we anticipate that this will play a similarly critical role with mass spectrometric analysis of small molecules in the future. In that perspective, we integrated `passatutto` into GNPS web platform to ensure that the community can readily search spectral libraries in high-throughput manner while reporting a significance of the annotation. We further envision that robust accuracy estimations, including FDR, will also enhance the analysis of spectral matches for in silico generated reference libraries or in silico annotations^4,36–41^, that are beginning to play important roles in brightening the dark matter of untargeted metabolomics.^4,42,43^

## Methods

### Spectral libraries and processing

We use three reference libraries for evaluating our FDR estimations: Agilent, MassBank and GNPS. The requirements for a MS/MS spectrum of a compound to be included in the analysis are that they had to a) have a SMILES or INCHI associated with it; b) to remove low resolution reference data, the exact parent mass must be within 10 ppm of the observed mass; c) the unfiltered MS/MS spectrum has to be available in the public domain. To ensure maximal homogeneity, we keep only d) spectra in positive ion mode, e) compounds below 1000 Da, and we discard f) spectra with less than 5 peaks with relative intensity above 2%. Spectra recorded at different collision energies are merged. In total, MS/MS spectra of 6,716 compounds (4,138 GNPS, 2,120 Agilent, 458 MassBank) fit these criteria. Most GNPS and all Agilent spectra were measured on Q-TOF instruments, all MassBank spectra were recorded on Orbitrap instruments. Not all peaks/signals in an MS/MS spectrum can be explained as fragment ions^34^, but we will stick with the term ‘fragment ion’ instead of ‘hypothetical fragment ion’, ‘peak’ or ‘signal’ for the sake of readability. Similarly, we will speak of the ‘mass’ of a fragment ion when we refer to the observed *m/z* value, and of its ‘intensity’ when we refer to the peak intensity.

### Noise filtering

For each target MS/MS spectrum, we calculate a fragmentation tree that annotates a subset of hypothetical fragment ions with molecular formulas^27,34,44^; only annotated fragment ions are kept, using the original peak intensities. We set mass deviation parameters 10 ppm (relative) and 2 mDa (absolute). This procedure is more sensitive than simply using a hard or soft intensity cutoff, as it ensures that fragment ions can be explained in principle by some sensible fragmentation cascade. After noise filtering, 104 spectra were empty or consisted only of the precursor ion peak, and were discarded.

### Software and creation of decoy databases

`passatutto` has been implemented as a Java v1.6 program. It reads and writes spectra in MassBank file format and fragmentation trees in the SIRIUS DOT file format. Source code is available from https://github.com/kaibioinfo/passatuto, Java executables (JAR files) are available from https://bio.informatik.uni-jena.de/passatutto/. `passatutto` contains modules for a) generating a decoy database, b) database searching in locally stored datasets, and c) estimating q-values either by means of the target-decoy approach or by empirical Bayes estimation. For generating a decoy database using the fragmentation tree-based method, SIRIUS can be used for the computation of fragmentation trees, which is available from https://bio.informatik.uni-jena.de/sirius/. We ran the software on an Intel XEON 6 Core E5-2630 at 2.30 GHz with 4 GB memory.

### FDR estimation

Given a decoy database, FDRs and q-values can be estimated using target-decoy competition^25^, separated target-decoy search^23^, or the more sophisticated mix-max approach^44^. Here, we use a simple separated target-decoy search^23^ where the proportion of incorrectly annotated spectra^33^ is estimated from the empirical Bayes distribution.

For the empirical Bayes approach^26,45^, we model database search scores as a two-component mixture of distributions representing true hits and false hits (true positive and false positive identifications). Scores of true hits are modeled using a mirrored Gamma distribution, a mirrored Gumbel distribution, or a mirrored Weibull distribution, whereas scores of false hits are modeled using a Gamma distribution. Both the actual distribution for true hits and the distributions’ parameters are chosen based on the observed data, where Expectation Maximization is used to simultaneously find the parameters of the mixture distribution. FDR and q-values are estimated using the average Posterior Error Probability of *all* hits with score above the threshold; the Posterior Error probability for a given score is estimated as the proportion of incorrect hits amongst *all* hits with this score.

### Quality assessment of FDR estimation

The two smaller datasets, MassBank and Agilent, are used as query spectra, whereas the larger GNPS dataset is searched in. The estimated p-value is the ratio of decoy hits with score above the threshold. As we know the true identity of all queries, we can calculate the true FDR (ratio of false hits among all hits) for any score threshold; by definition, the q-value of a hit is the smallest FDR for which it is reported.

### FDR based annotations for metabolomics

Passattuto produced decoy spectra for GNPS’s spectral library search workflows. These workflows were altered to enable FDR estimation utilizing these decoys to estimate provide q-values for all identifications in a search. The number of identifications were reported at 1% and 5% FDR for each of the 70 datasets analyzed in GNPS and were compared against the default scoring thresholds recommended at GNPS (0.7 cosine, 6 minimum matched peaks).

### The impact of scoring parameters that achieve 1% FDR

To evaluate how scoring settings such as cosine score and number of minimum ions to match affected the number of annotations with an FDR of 1%, we ran `passatutto` on the same 70 public projects but varies the number of minimum matched ions from 2 to 8. We then reported the number of data sets that achieved 1% FDR at each cosine value and minimum number of ions that matched. Finally, we reported the number of spectra matched for all of the different projects.

## Acknowledgements

We thank the Deutsche Forschungsgemeinschaft for supporting this work under grant BO 1910/16 and for a postdoctoral research fellowship to D.P. with grant number PE 2600/1-1. We thank the NIH for supporting this work under NIH number P41 GM103484 and the NIH grant on reuse of metabolomics data R03 CA211211.

### Code and data accessibility

Link to passatuto application and used spectral data:

https://bio.informatik.uni-jena.de/passatutto/

Github link to passatuto source code:

https://github.com/kaibioinfo/passatuto

Web based passatuto: http://gnps.ucsd.edu/ProteoSAFe/static/gnps-experimental.jsp

All metabolomics data used was from GNPS http://gnps.ucsd.edu

The accession numbers for the data sets used are MSV000078567, MSV000078584, MSV000078586, MSV000078589, MSV000078598, MSV000078603, MSV000078611, MSV000078612, MSV000078628, MSV000078649, MSV000078658, MSV000078670, MSV000078683, MSV000078708, MSV000078710, MSV000078711, MSV000078719, MSV000078726, MSV000078744, MSV000078805, MSV000078811, MSV000078812, MSV000078816, MSV000078832, MSV000078892, MSV000078903, MSV000078922, MSV000078936, MSV000078960, MSV000078993, MSV000079029, MSV000079040, MSV000079050, MSV000079069, MSV000079091, MSV000079104, MSV000079105, MSV000079146, MSV000079243, MSV000079329, MSV000079339, MSV000079341, MSV000079344, MSV000079356, MSV000079398, MSV000079416, MSV000079421, MSV000079447, MSV000079450, MSV000079558, MSV000079573, MSV000079581, MSV000079598, MSV000079651, MSV000079652, MSV000079679, MSV000079758, MSV000079760, MSV000079772, MSV000079773, MSV000079777, MSV000079778, MSV000079787, MSV000079808, MSV000079813, MSV000079825, MSV000079838, MSV000079888, MSV000079905, MSV000079907

